# Kappa Prophage Impacts *Vibrio cholerae* Fitness and is Restricted by BREX Phage Defense System

**DOI:** 10.64898/2026.10.16.759536

**Authors:** Andrea Rebolledo Viveros, David W Lazinski, Marzia Sultana, Munirul Alam, Andrew Camilli

## Abstract

*Vibrio cholerae*’s pathogenicity largely stems from the acquisition of cholera toxin genes via lysogenic conversion mediated by the CTX prophage. However, the strong association of other prophages with epidemic strains has not been linked to lysogenic conversion. During the first wave of the seventh cholera pandemic, the Kappa prophage was commonly found in clinical isolates and was later shown to confer a competitive fitness advantage in the environment and to accelerate evolution by stimulating horizontal gene transfer; however, its role in virulence remains unknown. Here, we report that the Kappa prophage confers a competitive fitness advantage *in vitro* and during intestinal tract infection in both Wave 1 and Wave 3 strains. This competitive fitness advantage is due to phage predation on neighboring cells, and not to lysogenic conversion. Furthermore, we show that loss of Kappa coincides with the acquisition of novel phage defense systems from SXT ICEs *Vch*Ban9 and *Vch*Ind5. Phage defense systems BREX and AbiC inhibited Kappa predation and restricted horizontal gene transfer, thereby reducing Kappa’s competitive fitness advantage.

**Importance:** As prophages are found in many bacterial genomes, understanding their roles in bacterial fitness advances our understanding of bacterial evolution and virulence drivers. The work presented here contributes to our understanding of the temperate phage-host arms race and highlights how the acquisition of phage defense systems can be a double-edged sword in *V. cholerae*, protecting it from phage predation yet limiting lysogenic conversion and evolution.

## Introduction

Bacteriophages (phages) are the most abundant biological entities and help shape microbial ecology, evolution, and pathogenicity. Among phages, integrated temperate phages (prophages) are found in over 70% of bacterial genomes [1] and have played a major role in bacterial evolution, primarily through horizontal gene transfer and lysogenic conversion (provision of pathogenicity genes, such as toxins). A notable example of this bacterial-phage evolutionary dynamic is *Vibrio cholerae*, the Gram-negative bacterium responsible for the severe diarrheal disease cholera. The pathogenicity of *V. cholerae* relies heavily on the CTXф prophage, which carries genes encoding cholera toxin (*ctxAB*). Integration of CTXф converts non-toxigenic *V. cholerae* strains into toxigenic ones capable of causing cholera outbreaks.

*V. cholerae* harbors other temperate phages [2], yet their roles in lysogenic conversion or other aspects of *V. cholerae* biology remain unknown. Kappa-type phages are the most characterized and are of particular interest because of their high prevalence in clinical isolates throughout the early seventh cholera pandemic and concerns about another lysogenic conversion [3–5]. Although no significant change in cholera toxin production or *V. cholerae* pathogenicity could be attributed to lysogenization by the prototype Kappa phage K139 [6], single-strain infections using the infant mouse model of colonization with *V. cholerae* K139 and K139 mutant lysogens provide supporting evidence for the Kappa phage *glo* gene as a possible virulence factor [5]. Yet a later study instead showed that *glo* participates in phage exclusion [4]. In an *in vitro* estuary-like microcosm, *V. cholerae* K139 lysogens were shown to out-compete non-lysogens through neighbor predation and promote horizontal gene transfer [7, 8]. Neighbor predation has also been shown to confer a competitive advantage to *Escherichia coli* lysogens in the intestinal tract of mice, albeit offset by a higher fitness cost due to a high prophage induction rate *in vivo* that results in host cell lysis [9].

For *V. cholerae*, the ecological advantage and possible lysogenic conversion support the maintenance of Kappa phage in the population. Yet, after decades of coexistence between Kappa phage and *V. cholerae*, the prevalence of Kappa lysogens declined in the early 2000s and has rarely been detected since [10]. Here, we examine the competitive fitness of K139 lysogens *in vitro* and in the infant mouse model of infection and explore the acquisition of phage defense systems throughout the 7^th^ pandemic to elucidate factors contributing to the decline of Kappa lysogens. We demonstrate that Kappa lysogens have a competitive fitness advantage *in vivo* due to neighbor predation, and the BREX phage defense system is necessary and sufficient to restrict Kappa predation and lysogeny. In addition, we identify compounds that trigger prophage induction that are relevant to intestinal infection.

## Results

### Bile salts induce Kappa prophage

To understand the role of Kappa prophage throughout the three Waves of the ongoing 7^th^ cholera pandemic and the reasoning for its disappearance, we first investigated Kappa prophage induction in Wave 1 and Wave 3 strains. We used the K139-like prophage from the El Tor Wave 1, 1978 clinical isolate E7946 to generate a spectinomycin-marked Kappa (hereafter referred to as Kappa-spec). We prepared Kappa-spec lysogens in the E7946 background and in two Wave 3 clinical strains isolated in 2021. Bile has been shown to play a role in *V. cholerae* virulence and biofilm production [11, 12] and to induce prophages in other bacteria that inhabit the human intestinal tract [13–15]. We sought to determine whether the disappearance of Kappa prophage was due to differences in induction by using semi-crude and individual bile salts as potential biological stressors and inducers of Kappa prophage. We determined that 0.05% chenodeoxycholate (CDC) or deoxycholate (DC) resulted in Kappa prophage induction in both Wave 1 and Wave 3 strains (**Fig. 1A**), which is consistent with temperate phage titers observed for other phages in *in vivo* studies [16, 17].

**Figure 1.**
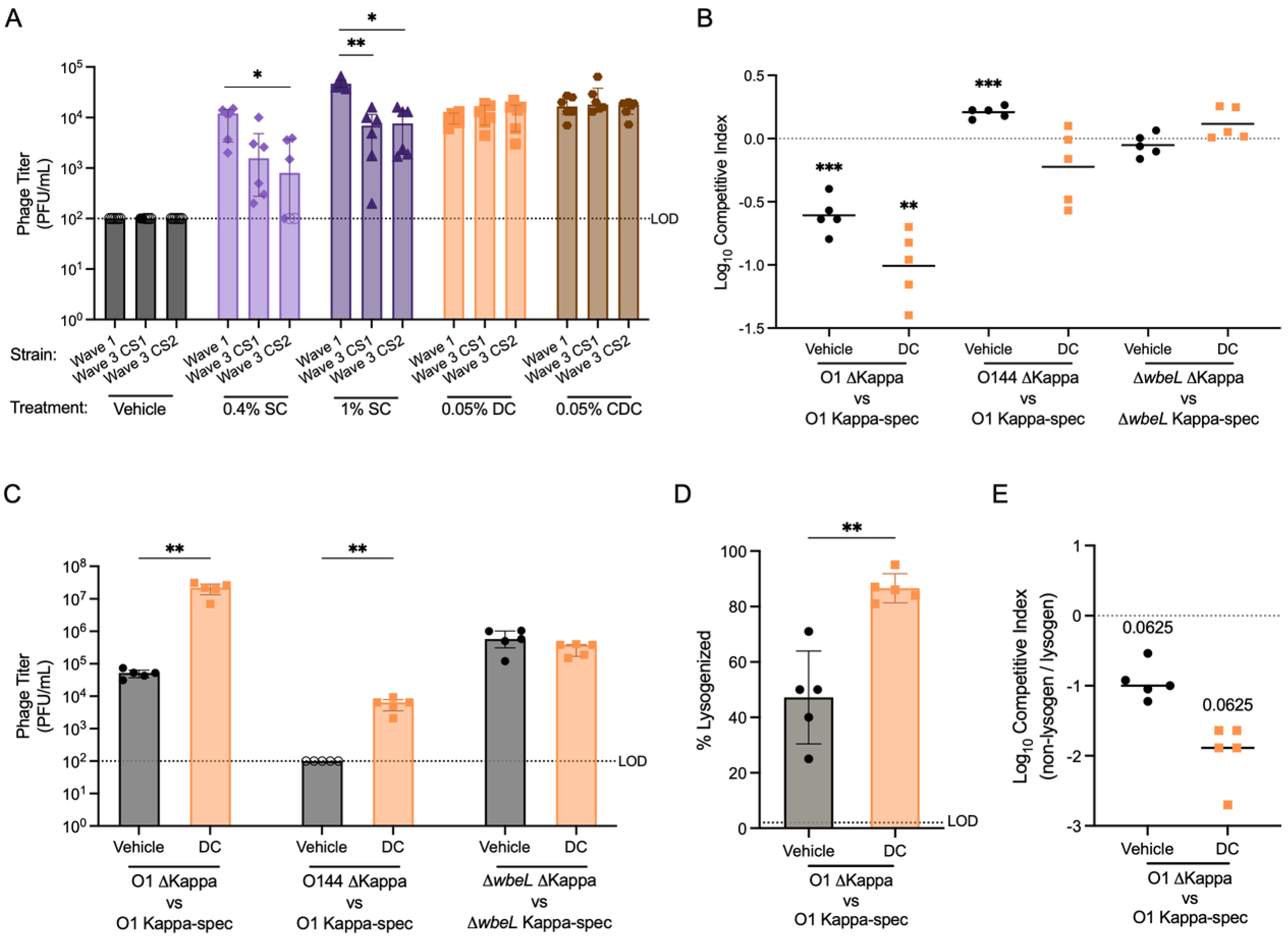
**Kappa lysogens outcompete non-lysogens in vitro through neighbor predation under inducing and non-inducing conditions in a Wave 1 background**. A) Phage titers from Kappa prophage induction using semi-crude (SC) or individual bile salts in Wave 1 E7946 and Wave 3 clinical strain 1 (CS1), and clinical strain 2 (CS2). B-D) Competition assays between Kappa-spec lysogens and non-lysogens in a Wave 1 E7946 background with either O1 antigen (Kappa receptor), O1 antigen genes swapped for those of O144, or the O1 synthesis gene *wbeL* deleted. B) Log10 competitive index under vehicle or 0.05% DC treatment. C) Phage titers from competition assays. D) Lysogeny percentage of non-lysogens at output. E) Log10 competitive index adjusted for lysogeny. Each symbol represents a biological replicate, with open symbols representing data points below the limit of detection (LOD). Shown are the median with interquartile range for A) and C), mean with SD for D), and mean for B) and median for E). Significant differences are represented by A) Kruskal-Wallis test with post hoc Dunn’s test, B) one-sample t test, C) Mann-Whitney test, D) Welch’s t test, and E) Wilcoxon signed-rank t test. p-values as follows: * p<0.05, ** p<0.01, and *** p<0.001.

### Kappa prophage provides a competitive fitness advantage through neighbor predation in vitro

Given our ability to induce Kappa-spec in both Wave 1 and 3 strains, it is unlikely that the lack of Kappa prophage induction can explain its decline during the 7^th^ pandemic. We next asked whether Kappa prophage provides a fitness advantage in Wave 1 strains by competing E7946 Kappa-spec with its non-lysogenic isogenic strain, with and without 0.05% DC. We found that Kappa-spec lysogens outcompete non-lysogens with and without induction, but to a greater extent with induction (**Fig. 1B**). In the vehicle-treated condition, there is some spontaneous Kappa-spec induction, followed by predation of non-lysogens and the resultant amplification of the phage, as phage titers in the vehicle-treated competition (**Fig. 1C**) are much higher than those in vehicle-treated lysogen alone (**Fig. 1A**).

If neighbor predation is occurring in the competition assays, we should observe some degree of lysogenization. Indeed, the percentage of initial non-lysogens that became lysogens ranged from 25-71% in the vehicle-treated competition and 81-95% in the 0.05% DC-treated competition (**Fig. 1D**). Due to the high rates of lysogenization occurring during the competitions and thus the transfer of spectinomycin resistance, the competitive indexes, calculated by blue-white colony screening, needed to be corrected to account for lysogenization of the competing strain. This is shown as the competitive index of total non-lysogens/lysogens (**Fig. 1E**). This correction yielded substantially higher out-competition values under vehicle- and DC-induced conditions than when lysogenization is not accounted for (**Fig. 1B**).

To address whether Kappa prophage provides a fitness advantage beyond neighbor predation, we generated competitor strains that cannot be preyed upon by eliminating the Kappa phage receptor, the lipopolysaccharide O1 antigen. Either the O1 antigen biosynthetic locus was replaced with that of the O144 antigen, or the O1 antigen gene *wbeL* was deleted, resulting in a rough mutant phenotype with only lipid A and core oligosaccharides produced [18]. The O144 antigen is chemically and structurally similar to the O1 antigen [19, 20], yet it cannot serve as a receptor for Kappa-spec, as no plaque-forming units (PFUs) were observed in this study. We performed competition assays with and without 0.05% DC and found that, in the absence of neighbor predation, Kappa-spec lysogens no longer have a competitive fitness advantage with either competitor strain (**Fig. 1B**). Consistent with the lack of neighbor predation, phage titers were below the limit of detection for the vehicle-treated O144 antigen competition (**Fig. 1C**). In contrast, and unexpectedly, there were high phage titers in vehicle- and DC-induced competitions with the Δ*wbeL* strain (**Fig. 1C**). This result is likely due to the Kronos effect, in which phage progeny are lost through reinfection of lysogens [21], and it is supported by Kappa-spec adsorption to the various strains **(Fig.S1)**. In this case, Kappa-spec fails to productively adsorb to and infect the rough mutant, whereas it adsorbs to the O1 lysogens in the O144 vs O1 competition, resulting in the inactivation of Kappa phage.

### Kappa prophage provides a competitive fitness advantage in vivo

We examined whether the competitive fitness advantage observed *in vitro* is evident *in vivo* using an infant mouse model of small-intestinal colonization. In both Wave 1 and 3 backgrounds, the lysogenic strains have a competitive advantage *in vivo* (**Fig. 2A**). Consistent with neighbor predation as the mechanism, 40-50% of the competing strains were lysogenized (**Fig. 2B**), and phage titers were detectable at levels above 10^4^ PFU/mL in the small intestine homogenates(**Fig. 2C**). To confirm that lysogenization was occurring, we deleted the attB site in the Wave 1 non-lysogen strain and showed that lysogenization was largely abolished (**Fig. 2B**).

**Figure 2.**
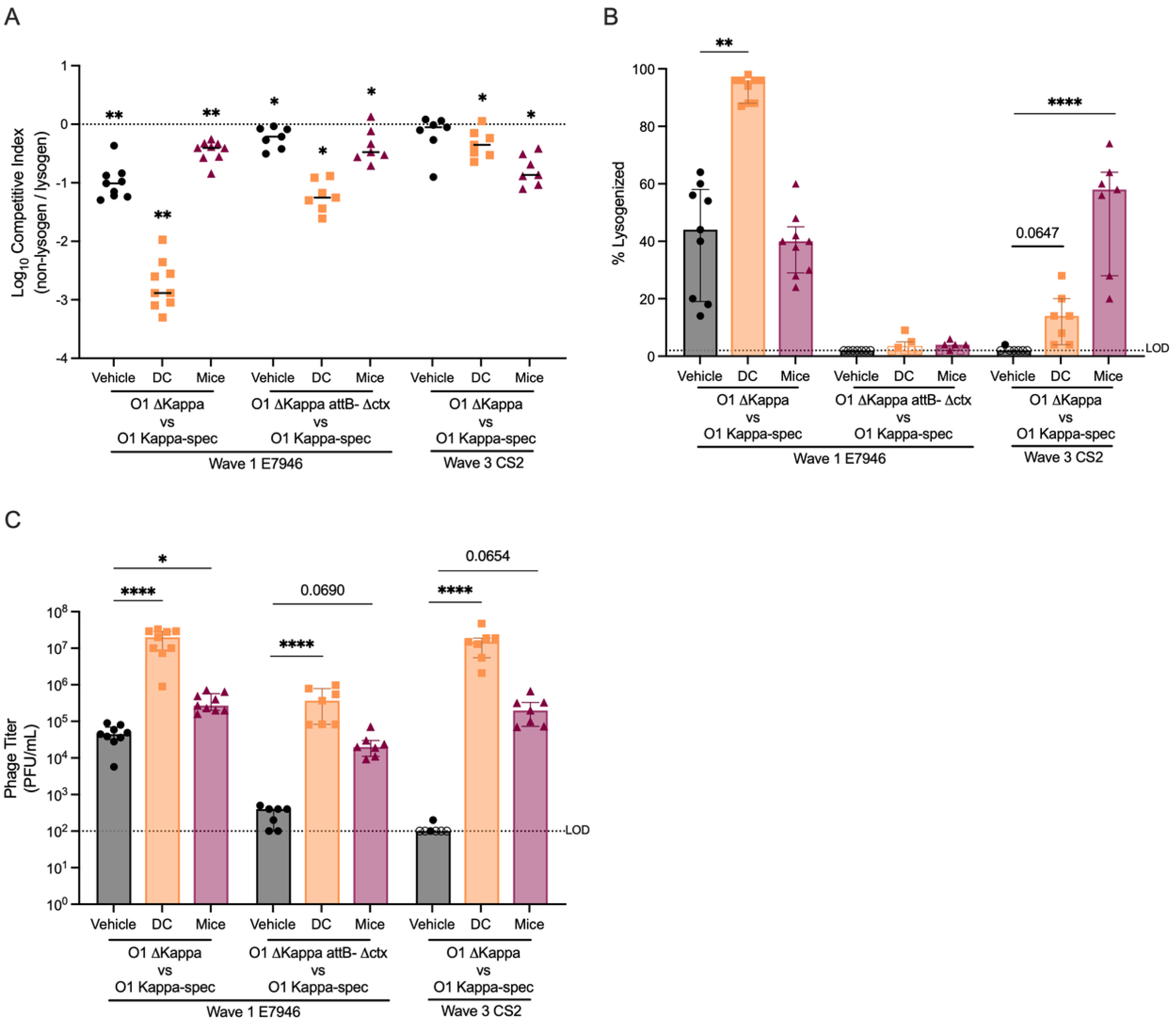
**Kappa lysogens outcompete non-lysogens in both Wave 1 and Wave 3 strains**. A) Log_10_ competitive index adjusted for lysogeny in Wave 1 and Wave 3 backgrounds under different treatments. B) Percent of non-lysogens that were Kappa-spec lysogenized. C) Phage titers from corresponding competitions. Each symbol represents a biological replicate, with open symbols representing values below the limit of detection; hence, the limit of detection value was assigned. The median is shown for all, with the interquartile range for B-C). Significance differences represented are from A) Wilcoxon Signed Rank test, and B-C) Kruskal-Wallis test post-hoc Dunn’s test. p-values are represented as: * p<0.05, ** p<0.01, *** p<0.001, and **** p<0.0001.

In contrast to the Wave 1 K139 lysogen, the Wave 3 lysogen showed no detectable spontaneous prophage induction *in vitro* in the vehicle-treated competitions, but did in the presence of DC (**Fig. 2A-C**). In an infant mouse model, the Wave 3 strain Kappa-lysogen outcompeted the non-lysogen at a ratio similar to that seen for the Wave 1 strain, resulting in similar lysogeny and phage titers. This is consistent with preferential prophage induction *in vivo* compared to 0.05% DC *in vitro.* Notably, the Wave 3 *in vitro* DC competitions yielded a mix of turbid and clear plaques, with the latter representing a small percentage of the total plaques. Isolation of these clear plaques and K139 whole-genome sequencing confirmed that they were spontaneous lytic mutants of Kappa-spec harboring mutations in the *cI* and *cII* repressor genes. Taken together, our results suggest that the decline in Kappa lysogens during the 7^th^ pandemic was not due to reduced induction or competitive fitness but rather other factors, possibly the acquisition of phage defense systems.

### The acquisition of the BREX phage defense system coincides with the disappearance of Kappa

Bioinformatic analysis of the genomes of 61 El Tor O1, seventh pandemic strains isolated between the years 1961 and 2021 for the presence of Kappa prophage and phage defense systems reveals the acquisition of phage defense systems AbiE and Bacteriophage Exclusion (BREX), around 1993, and AbiC around 2002, which coincides with the disappearance of Kappa prophage isolates (**Fig. 3**). These three systems are all located within the mobile SXT integrative and conjugative element (ICE) *Vch*Ind5. BREX defense systems are widespread across microbial genomes, protecting in a methylation-dependent manner, although the underlying restriction mechanism remains unclear [22–24]. Both AbiC and AbiE were characterized as abortive infection systems in *Lactococcus lactis* [25]. AbiE has been further characterized and reported to be a two-gene toxin-antitoxin system [26]. The AbiC mechanism remains unclear, but it has been shown to reduce capsid production and restrict plaque formation in different organisms [27, 28]. Of note, the phage defense predictor program DefenseFinder annotated AbiC as AbiJ, whereas the alternative program PADLOC [29] identified it as AbiC. AbiC and AbiJ share homology.

**Figure 3.**
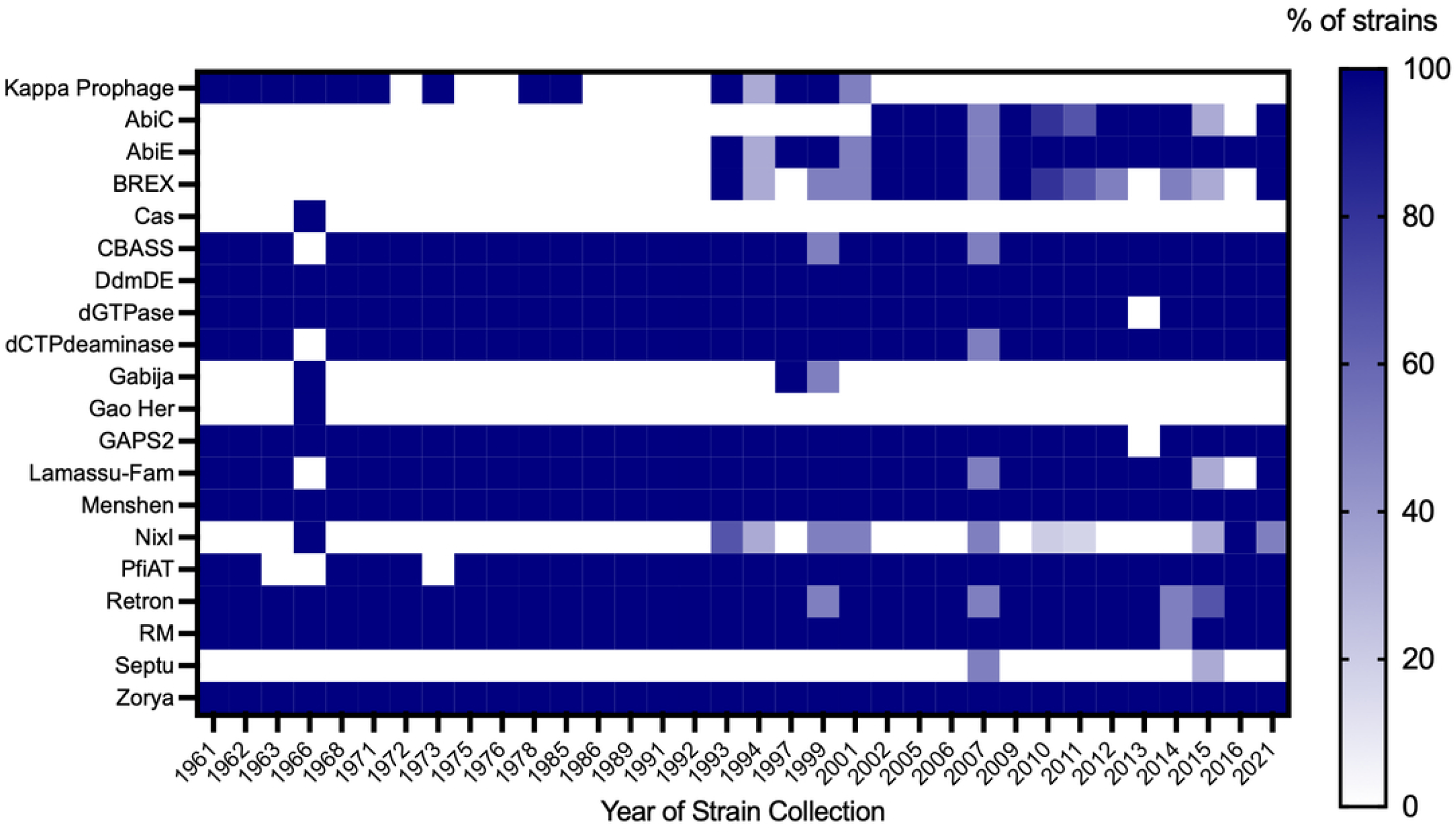
**Acquisition of novel phage defense systems in *V. cholerae* coincides with the decline of the Kappa prophage**. Heat map illustrating the percentage of strains in each year containing Kappa prophage and putative phage defense systems, based on analysis of 61 *V. cholerae* El Tor strain genomes spanning the years 1961 to 2021.

Of the three, only BREX is known to have activity against vibriophages. To test whether acquiring any of these phage defense systems contributed to the disappearance of Kappa phage, we introduced arabinose-inducible chromosomal insertions of each system into a Wave 1 strain that naturally lacks all three, and assayed Kappa-spec plaque formation and lysogenization. We found that induction of BREX reduced Kappa-spec efficiency of plating (EOP) by approximately six orders of magnitude, whereas AbiC and AbiE had no effect (**Fig. 4A**). The EOP of the lytic Kappa-spec mutant was reduced by four orders of magnitude. Notably, the lytic Kappa-spec mutant plaque-forming units did not dilute as expected, suggesting that multiple lytic phages infecting a single cell can bypass BREX restriction, albeit inefficiently. Consistent with restriction of wild-type Kappa-spec infection, BREX eliminated Kappa-spec lysogeny (**Fig. 4B**).

**Figure 4.**
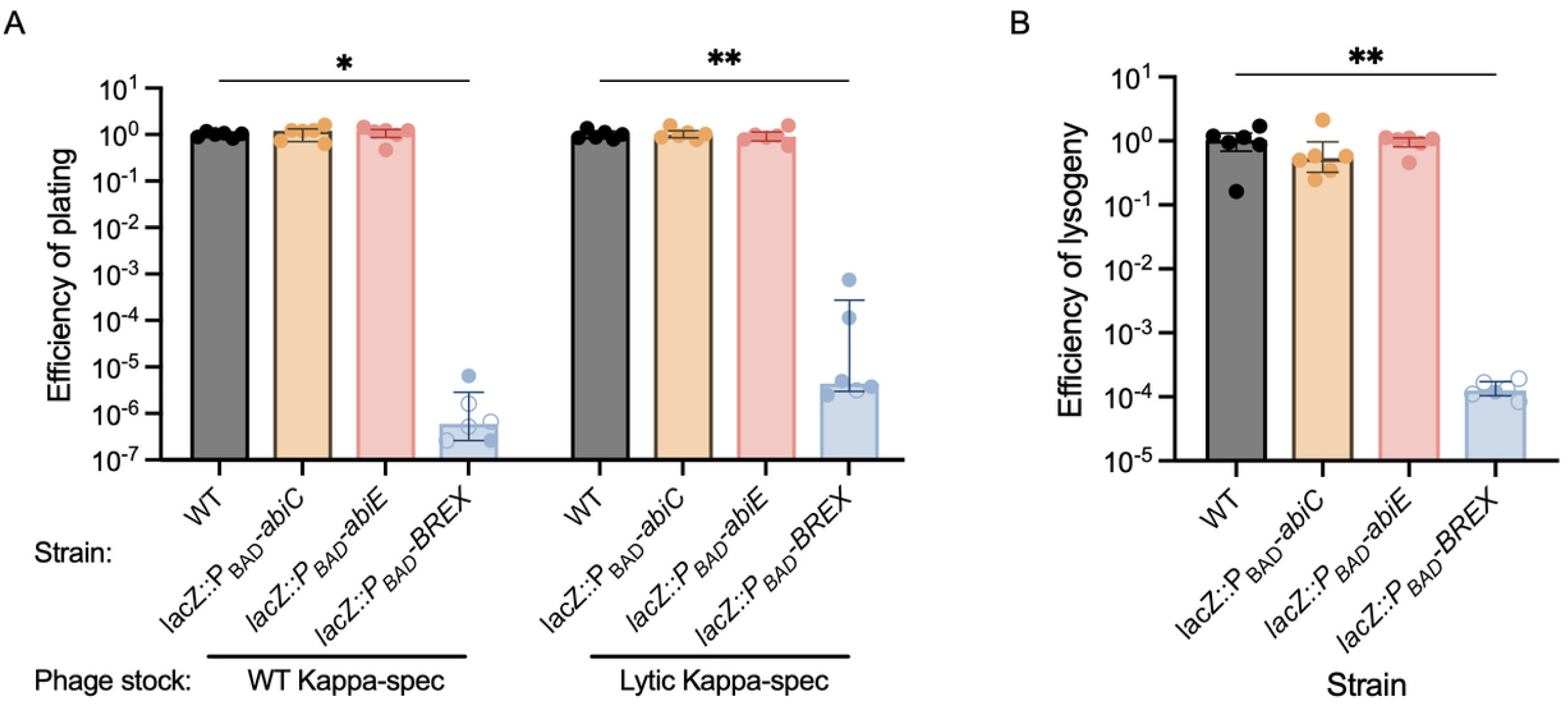
**Overexpression of BREX inhibits Kappa’s ability to plaque and lysogenize**. A) WT Kappa-spec or lytic mutant Kappa-spec efficiency of plating on WT, *abiC*, *abiE*, and BREX arabinose-inducible strains. B) WT Kappa-spec efficiency of lysogeny on WT, *abiC*, *abiE*, and BREX arabinose-inducible strains. Median with the interquartile range is shown. Each circle represents a biological replicate, with open circles representing values below the limit of detection; hence, the limit of detection value was assigned. Significant differences represented from the Kruskal-Wallis test post-hoc Dunn’s test as: * p<0.05 and ** p<0.01.

### BREX is necessary and sufficient for Kappa phage resistance

To determine whether restriction of Kappa-spec was an artifact of BREX overexpression, we generated a single-copy chromosomal insertion of BREX under its native promoter in the Wave 1 strain background. We then measured the Kappa-spec EOP and lysogenization. In a BREX+ strain, the EOP was reduced by more than five orders of magnitude (**Fig. 5A**), and lysogeny was below the limit of detection (**Fig. 5B**). Consistent with BREX functioning via methylation, if the phage stock is generated in a BREX+ strain, this inhibition is void (**Fig. 5A-B**). To confirm this mechanism, we generated a catalytically inactive mutant of the BREX methyltransferase BrxX (G290Y). When the phage stock is generated in this mutant background, restriction of EOP and lysogeny by wild-type BREX is restored (**Fig. 5A-B**). These data support a model in which BREX methylation of the Kappa phage genome allows it to bypass BREX restriction and are consistent with results observed with *E. coli* λ phage [24]. As both Wave 3 clinical strains (CS1 and CS2) used in this study contained BREX and required a high MOI for initial generation of Kappa-spec lysogens, we evaluated whether BREX alone was necessary to impose this restriction. Deletion of the BREX system from CS2 increased plating and lysogeny efficiencies (**Fig. 5C-D**) to a similar magnitude as observed for the Wave 1 strain. Thus, BREX is both necessary and sufficient for Kappa phage inhibition. However, the presence of both the Kappa prophage and the BREX system within some strains in 1993 and 2001 raised the question of what differed between these strains and later ones. All those strains contained the SXT *Vch*Ban9, while strains from 2002 onward contained the SXT *Vch*Ind5, whose type I BREX systems have been shown to vary and share little homology [30]. Thus, we compared the methylation effects of the two BREX systems, *Vch*Ban9 and *Vch*Ind5, using Kappa phage EOP and found that both systems restrict Kappa, but methylation of Kappa by one BREX system did not allow it to bypass the other (**Fig. 5E**). This exemplifies how variability within one organism’s SXTs, specifically within BREX systems and their different recognition sequence motifs, can play a role in preventing consistent phage predation and maintenance within the population.

**Figure 5.**
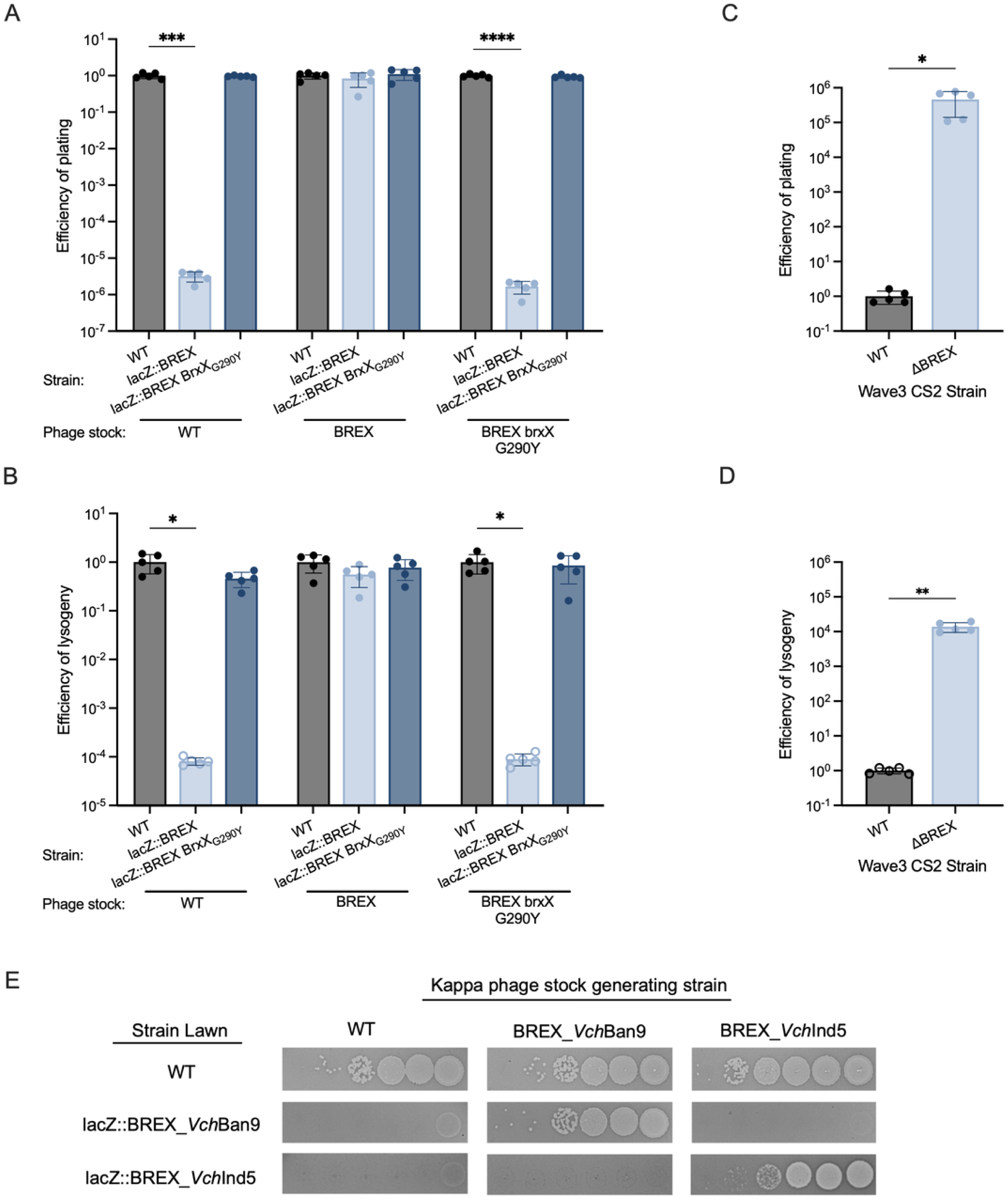
BREX is sufficient and necessary to inhibit Kappa prophage, yet methylation allows it to bypass inhibition. A) Efficiency of plating and B) efficiency of lysogeny of Kappa-spec stock generated in Wave 1 WT, BREX, and BREX BrxX G290Y on corresponding strains. WT Kappa-spec C) efficiency of plating and D) efficiency of lysogeny on Wave 3 WT and ΔBREX strains. E) Comparison of Kappa phage plaquing efficiency based on different *V. cholerae* STX BREX systems. Bars represent the mean with SD. Each circle represents a biological replicate, with open circles representing values below the limit of detection; hence, the limit of detection value was assigned. Significant differences represented are from A-B) Welch’s ANOVA post-hoc Dunnett’s test, and C-D) Welch’s t test with p-values as follows: * p<0.05, ** p<0.01, *** p<0.001, and **** p<0.0001.

### Competitive fitness advantage of Kappa lysogeny is outweighed by BREX protection against other vibriophages

Given the magnitude of Kappa phage inhibition by BREX, this raised the question of whether the acquisition of BREX by a non-lysogen would eliminate the competitive advantage that Kappa lysogens normally exhibit through neighbor predation. To begin to address this, we competed a Wave 1 Kappa-spec lysogen with a Wave 1 BREX+ non-lysogen strain *in vitro*. The presence of BREX in the non-lysogen abrogated the competitive fitness normally observed for the Kappa-spec lysogen, instead resulting in a slight loss of competitive fitness in vehicle-treated and a substantial loss of competitive fitness for DC-treated competitions (**Fig. 6A**; compare to Fig. 1B). The large loss of competitive fitness in the DC-treated competition presumably resulted from prophage activation and lysis of a substantial portion of the lysogen strain. Consistent with the lack of neighbor predation, the phage titer from the competition assays (**Fig. 6B**) matches that observed when phage predation was inhibited by competing against the O144 strain (**Fig. 1C**). These results are consistent with the hypothesis that the acquisition of BREX contributed to the loss of Kappa lysogens from the population, tilting the scale in favor of the BREX+ non-lysogens.

**Figure 6.**
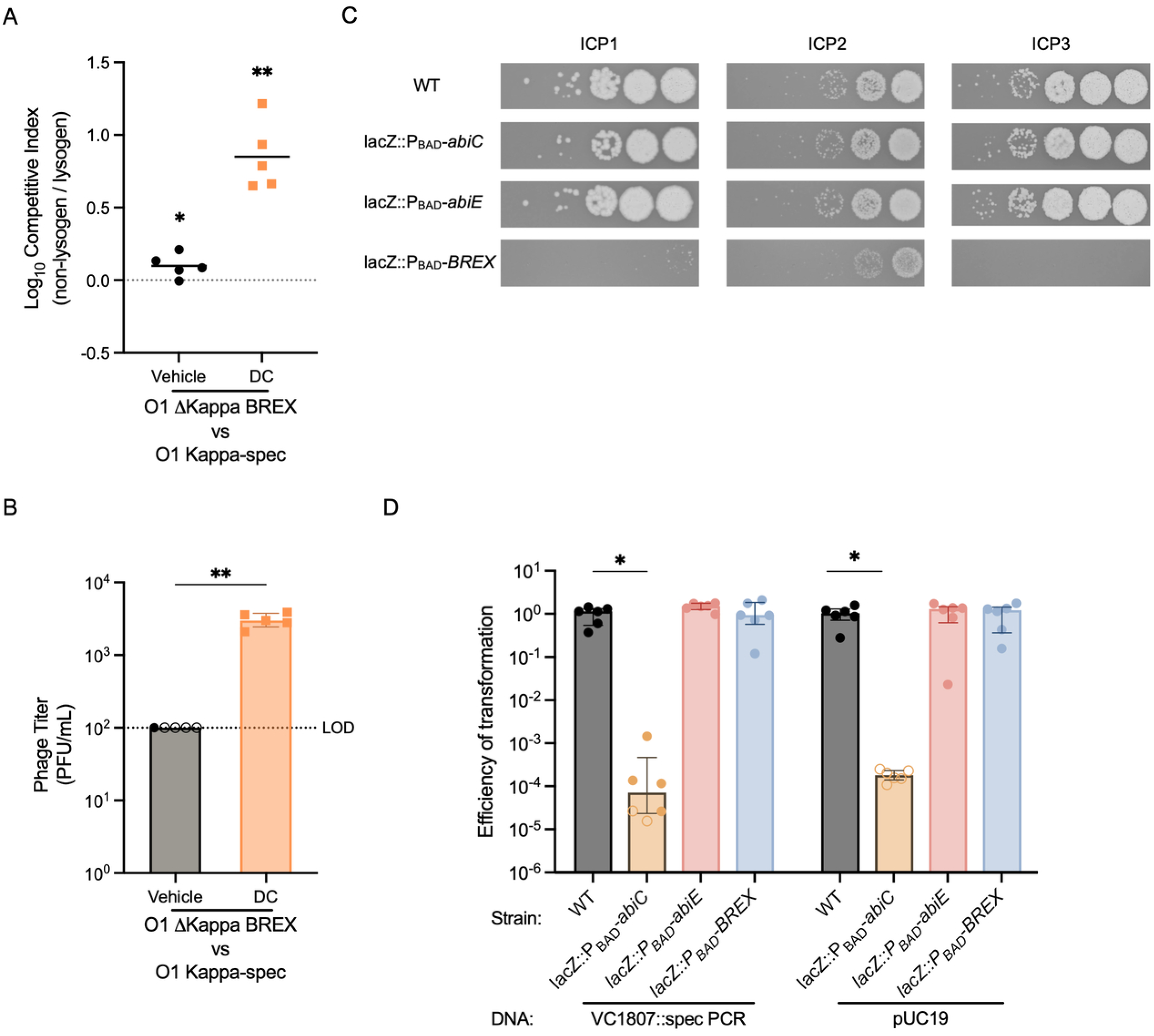
BREX and AbiC reduce Kappa lysogen competitive advantage. A) Log10 competitive fitness and B) phage titer from Wave 1 BREX non-lysogenic strain vs Wave 1 WT Kappa-spec lysogen in vitro competition. C) Plaquing efficiency of ICP1-3 on WT or arabinose-inducible strains of abiC, abiE, and BREX. D) Transformation efficiency of PCR fragment VC1807::SpecR or pUC19 on WT or arabinose-inducible strains of *abiC*, *abiE*, and BREX. Mean is shown for A), and median with interquartile range for B) and C). Each circle represents a biological replicate, with open circles representing values below the limit of detection; hence, the limit of detection value was assigned. Significant differences represented are from A) One sample t test, B) Mann-Whitney test, and D) Kruskal-Wallis test post-hoc Dunn’s test with p-values as follows: * p<0.05, and ** p<0.01.

However, Kappa was not the sole vibriophage present during the transition to BREX+ strains. ICP1, ICP2, and ICP3 are virulent phages consistently isolated from cholera patients’ stool samples [31], with ICP1 being a persistent *V. cholerae* predator in both estuaries and cholera patients, thereby driving ongoing defense and counter-defense evolution between ICP1 and *V. cholerae* [32, 33]. Therefore, we asked whether the induced expression of AbiC, AbiE, or BREX affected plaque formation by each of these three phages. BREX, but not AbiC or AbiE, greatly reduced the EOP of ICP1 and ICP3, and reduced that of ICP2 modestly (**Fig. 6C**). In addition, plaque sizes were smaller for ICP1 and ICP2. The reduction in the EOP for ICP3 upon overexpression of BREX was much greater than the reported 10-fold reduction under native BREX expression [30], indicating that expression level is limiting in the latter.

### AbiC inhibits the natural transformation of V. cholerae

Finally, because some phage defense systems have been shown to inhibit horizontal DNA transfer processes such as plasmid stability and conjugation [34, 35], we wondered whether AbiC, AbiE, or BREX might inhibit natural transformation in *V. cholerae*, an evolutionary process shown to occur during Kappa-mediated neighbor predation in an estuary microcosm [7]. Overexpression of AbiC, but not AbiE or BREX, resulted in more than three orders of magnitude reduction of natural transformation efficiency regardless of DNA type, linear or plasmid (**Fig. 6D**). The lack of BREX activity could not be explained by the absence of the recognition motif, as both DNA samples contained the published recognition motif for *Vch*Ind5 [30]. Overall, our data show that of the three putative systems tested, only BREX exhibits detectable phage defense activity under the conditions tested, whereas AbiC inhibits natural transformation.

## Discussion

Our results reveal that *V. cholerae* Kappa phage lysogens confer a competitive fitness advantage *in vitro* and in an infant mouse infection model through neighbor predation, but not through lysogenic conversion. Additionally, Kappa provides superinfection immunity to its host. Thus, the loss of Kappa during the seventh pandemic appears contradictory. Our bioinformatic analysis and experimental results suggest that the acquisition of BREX contributed to the decline in Kappa lysogens. One explanation, supported by our data, is that BREX inhibition of prominent virulent vibriophages ICP1, ICP2, and ICP3 is more advantageous to *V. cholerae* than the neighbor predation and superinfection immunity provided by the Kappa prophage. Aside from encoding BREX, the SXT ICE also provides additional benefits, such as conferring antibiotic resistance, making it, as a whole, more advantageous than the Kappa lysogen fitness advantage alone. Although we showed that induction of a Kappa lysogen from a BREX+ strain is sufficient to bypass BREX restriction in subsequent BREX+ hosts, this phenomenon appears to have been insufficient to prevent loss of Kappa. This could be in part attributed to variation in sequence recognition motifs among BREX systems within two of *V. cholerae* SXT ICEs, which does not allow for transferable resistance.

It is possible that other phage defense systems acted in concert with BREX to reduce Kappa in the population. We tested two such systems, AbiC and AbiE, deemed likely based on bioinformatics, but neither restricts Kappa. However, as *V. cholerae* naturally inhabits estuary environments, the effect of all these defense systems in an estuary environment should also be explored.

It remains unclear why Kappa lysogeny was so low in *V. cholerae* Wave 3 strains under *in vitro* conditions compared to Wave 1, yet was equally high in the infant mouse model of intestinal infection. This phenotype, along with the isolated lytic Kappa mutants observed after *in vitro* competition, suggests that BREX might be more active *in vitro* than in the mouse model. However, it is possible that Wave 3 strains possess additional phage defense systems that limit Kappa lysogeny, and lytic mutants might more easily bypass these restrictions.

Although we focused on phage defense systems as potential factors in Kappa’s decline, we would be remiss not to consider other causes. For example, in the classical *V. cholerae* biotype, the Kappa-type phages VcA-1 and VcA-2 produce a mixture of abnormal virus particles, a phenomenon also observed in an El Tor strain [36, 37]. These defective particles include tailless heads, headless tails, and heads with various tail lengths and shapes. Therefore, defective Kappa phage particles may have contributed to its decline, and the overall frequency of these aberrant particles should be investigated. Moreover, since Kappa prophages have been found in the genomes of non-O1 *V. cholerae*, environmental *V. cholerae* may still serve as a reservoir of Kappa prophages and allow for predation in non-BREX strains. This is supported by the detection of Kappa lysogens in clinical isolates from three cholera patients in 2013 [10], with one patient harboring a mixed population of Kappa lysogens and non-lysogens, highlighting *in vivo* competition. Our analysis of these clinical strains indicates the presence of an alternative SXT ICE, *Vch*Ind6, that lacks both BREX and AbiC. Thus, whether Kappa-type phages are truly absent from *V. cholerae* populations remains to be confirmed.

Finally, we showed that AbiC, but not AbiE or BREX, inhibits natural transformation in *V. cholerae* within an estuarine microcosm. Given that *V. cholerae* has hotspots of phage defense genes scattered throughout its mobile genetic elements, and that within these elements different defense systems, such as DdmDE and AbiC, have been present for decades, this raises questions about the extent of their role in restricting horizontal gene transfer by natural transformation or other processes such as conjugation. In our 61-strain collection, we observe that the presence of AbiC and BREX fluctuates, whereas DdmDE remains stable. Therefore, a large-scale population analysis of phage defense systems, from both patient samples and environmental reservoirs, could help elucidate the factors driving these selections. Our study further underscores the interaction between phage defense systems within *V. cholerae* mobile genetic elements and their roles in fitness, phage resistance, and the ongoing evolution of *V. cholerae*.

## Methods

### Bacterial strains and growth conditions

Bacterial strains were cultured in Miller lysogeny broth (LB) with aeration or on LB agar at 37°C. When appropriate media was supplemented with 40 µg/mL Xgal, 50 µg/mL carbenicillin (carb), 50 µg/mL kanamycin (kan), 100 µg/mL streptomycin (strep), 100 µg/mL spectinomycin (spec). Strains used are detailed in Supplemental Table 1.

### Generation of mutant strains

Antibiotic resistance cassettes were used to replace genes by stitching homology arms to the cassettes via splicing-by-overlap-extension (SOE) PCR or NEB HiFi. *V. cholerae* was transformed by either natural transformation [38] or chitin-independent transformation [39], followed by plating on selective LB agar plates. Strains were whole-genome sequenced on the Illumina NextSeq or NovaSeq platforms at the Tufts University Core Facility (http://tucf-genomics.tufts.edu) or on the Nanopore platform.

### Generation of Kappa-spec lysogenic strains

Kappa-spec was generated in Wave 1 strains by adding a spectinomycin resistance cassette after the 3’ terminal end of gene *kp45* in the Kappa prophage genome of strains E7946 and E7946 Δ*wbeL* by natural transformation with a PCR product, followed by selection on LB spec plates. All other Wave 1 E7946 and Wave 3 Kappa-spec lysogenic strains were generated by infecting strains at MOI=1 or MOI=10 with a Kappa-spec phage stock from E7946 Δ*wbeL* and selecting on LB spec plates. E7946 Δ*wbeL* was routinely used to prep Kappa-spec phage stocks because it generates much higher titer lysates than the wild-type host.

### Kappa-spec bile induction assays

Kappa-spec lysogen cultures were back-diluted into 2 mL LB cultures at an OD600 of 0.01, supplemented with semi-crude bile, sodium deoxycholate, sodium chenodeoxycholate, or vehicle (water), and incubated overnight (18-20 hours). Cells were pelleted by centrifugation at 5,000 RCF for 3 minutes, and the supernatant was incubated with 5% chloroform at room temperature (RT) for 1 hour. The aqueous layer was used to enumerate the phage titer on soft agar lawns of E7946 ΔKappa by spot-plating 10 µL of serial dilutions.

### Competition assays

Overnight cultures were back-diluted and grown to mid-exponential growth phase (OD600 0.3-0.6). Each pair of competing strains, one LacZ^+^ and the other LacZ^-^, was mixed at a 1:1 ratio at an OD600 of 0.01 in LB and split into two treatment groups: 1) infant mice infection, and 2) 2mL *in vitro* cultures supplemented with 0.05% DC or an equal volume of water (vehicle). The input was serially diluted in LB, and 10 µL was dribbled in duplicate on LB Xgal strep plates (all strains used in this study were streptomycin-resistant) to determine the ratio of initial lysogen and non-lysogen based on blue and white colonies. Competitions were allowed to proceed for 20-24 hours.

*In vitro* output was processed by centrifuging cells at 5,000 RCF for 3 minutes, then resuspending them in fresh LB. The supernatant was incubated with 5% chloroform for 1 hour at RT, and the aqueous layer was used to enumerate the phage titer on lawns of AC7800, a Kan-resistance derivative of E7946 ΔKappa. Output cells were serially diluted, and 10 µL was dribbled in duplicate on LB Xgal strep and LB kan plates to determine the ratio of non-lysogen to lysogen and to distinguish the percentage of non-lysogen that were lysogenized. Initial non-lysogen strains contained a Kan-resistance cassette. To determine the percentage of non-lysogens that were lysogenized, 50 colony-forming units (CFUs) from each LB kan treatment plate were re-streaked on LB spec plates, and the percent lysogenized was calculated as the number of CFU in LB spec divided by 50, with the result multiplied by 100.

The competitive index (CI) was calculated as the (output ratio of non-lysogen/lysogen)/(input ratio of non-lysogen/lysogen) from blue-white colony screening. To account for lysogenization of the competing non-lysogen strain, the total number of non-lysogens that were lysogenized was estimated as X = CFU of non-lysogens in the output multiplied by the percent lysogenized. Then we calculated the adjusted non-lysogen and lysogen output population by subtracting X from the non-lysogen count and adding X to the lysogen count. The adjusted CI was calculated as the (adjusted output ratio of non-lysogen/lysogen) / (input ratio of non-lysogen/lysogen).

### Infant mice infection

All animal experiments were in accordance with the rules of the Tufts Comparative Medicine Services and the Institutional Animal Care and Use Committee. Four- and five-day-old CD-1 infant mice (Charles River Laboratories) were infected. Each group of mice was comprised of animals from at least two different litters. Both male and female animals were used in this study. The number of mice used in each condition in each experiment is indicated by the number of symbols shown in the figures. Infant mice were orogastrically inoculated with 10^5^ CFU of *V. cholerae* diluted in LB. Mice were killed 20-24 h post-infection. Small intestines were dissected and homogenized in PBS with 20% glycerol. *V. cholerae* CFU and Kappa-spec PFU were enumerated as per *in vitro* processing with the following modification for phage titer: 500 µL of homogenate was centrifuged at 10,000 RCF for 1.5 min, and the supernatant was incubated with 5% chloroform.

### Kappa prophage and defense system bioinformatic analysis

A search at the US National Center for Biotechnology Information was conducted for *V. cholerae* O1 El Tor strains isolated between 1961 and 2016, plus two previously reported Wave 3 clinical strains isolated in 2021 [40]. We sequenced the genomes of the two strains, 1 (DVC-512; CS1) and 2 (DVC-546; CS2), on the Novaseq platform (Illumina Inc.) at the TUCF, and by Nanopore sequencing (Plasmidsaurus.com). The presence of Kappa prophage was determined through a BLAST search of the Kappa phage NCBI Reference Sequence NC_010275 against the *V. cholerae* genomes. Only full genomes were found.

*V. cholerae* genomes were run through the computer program DefenseFinder [41] and predicted phage defense systems were annotated. For those years in which multiple strains were analyzed, the average percent presence of the Kappa prophage and each defense system was calculated and reported in the heatmap (Figure 3). Complete data with accession numbers is compiled in Supplemental Table 2.

### Generation of high-titer Kappa-spec stocks

Kappa-spec lysogens were generated in Wave 1 Δ*wbeL* E7946 WT, BREX+, and BREX+ BrxX G290Y backgrounds. Cultures were grown overnight and back-diluted 1/500 in 1 L LB. Then shaken at 37°C for 10-18 hours. For lytic Kappa-spec mutant, a 500 mL culture of AC6145 at OD=0.01 was infected at MOI=0.1 and shaken at 37°C for 5 hours. Cells were pelleted at 7,500 RCF for 15 min at 4°C and the supernatant was filtered through 0.22 µM PES membranes. Polyethylene glycol MW 8000 powder was dissolved in phage supernatant to a final concentration of 10% w/v. The phages were pelleted by centrifugation at 10,000 RCF for 45 min at 4 °C, and resuspended in 1 mL Sodium Chloride-Tris-EDTA (STE) buffer. Resuspended phage was filtered through a 0.22 µM PES syringe filter and stored at 4°C. Phage titer was enumerated on E7946. Phage genomes were sequenced on Nextseq platform (Illumina Inc.) at the TUCF.

### Efficiency of plating

Overnight cultures were back-diluted and grown to mid-exponential growth phase (OD600 0.4-0.6) at 37 °C with aeration. When called for, all strains, including the wild-type control, were incubated with 0.1% arabinose throughout. Lawns were prepared by combining 9 mL of 0.5% top agar with 1.62 × 10^8^ cells in 100-mm petri plates. Phage stocks were serially diluted in LB, and 10 µL dribbled onto the corresponding lawns. Plates were incubated at 37°C overnight. EOP was calculated as PFUs on mutant strain lawn / average number of PFUs on the WT strain lawn. For general visualization,10 µL of phage dilutions were spotted.

### Efficiency of lysogeny

Overnight cultures were back-diluted and grown to mid-exponential growth phase (OD600 0.4-0.6) at 37 °C with aeration. When needed, all strains, including the wild-type control, were incubated with 0.1% arabinose until selective plating. 5 x10^8^ cells were resuspended in 250 µL LB in microfuge tubes and phage stock added at an MOI=0.01. The mixture was incubated rolling at 37 °C for 10 minutes, followed by washing with 250 µL LB and resuspending in 100 µL LB. Cells were then incubated for an additional 15 minutes, rolling at 37 °C, before serially diluting in LB, and dribbling 10 µL onto LB strep and LB spec plates. The percent lysogenized was calculated as (number of CFUs on spec plates/number of CFUs on strep plates) × 100. The efficiency of lysogeny was calculated as the percentage of lysogenized of mutants/the average percentage of wild-type lysogenized.

### Efficiency of transformation

Overnight cultures were back-diluted and grown to mid-exponential growth phase rolling at 30 °C with 0.1% arabinose. Arabinose induction was maintained until DNA was added. Cells were washed and resuspended in 0.7% Instant Ocean (Instantocean.com) to an OD600 of 0.25, and 700 µL was added to 80 mg of chitin in a 2 mL microfuge tube. Natural competence was induced by static incubation with the microfuge lid closed for 22 hours at 30 °C. The culture supernatant was then carefully removed without agitating the chitin biofilms, and an equal volume of fresh 0.7% Instant Ocean was added with 250 ng of the pUC19 or VC1807::specR PCR product, and the mixture was incubated overnight at 30 °C under static conditions. The VC1807::specR PCR product, with its 1.8 and 2 kb arms of homology, was generated by PCR with primers 5’-GACGATTCTCATGCCGCTATTGC-3’ and 5’-GTTCTGCTCGATACTACATTGTGAG -3’, using 50 ng of genomic DNA of spectinomycin-resistant E7946 VC1807::SpecR derivative AC5136 as template. After 22 hours, the microfuge tubes were vigorously vortexed for 1 minute to dislodge cells from the chitin, and 500 µL was transferred to 1 mL LB and allowed to recover at 37 °C, rolling for 1 hour. The recovered cells were serially diluted in LB and assayed by dribbling 10 µL for total CFUs on LB strep agar plates and for transformants on LB carb or LB spec agar plates. The percent transformed was calculated by (number of CFUs on LB carb or LB spec plates/number of CFUs on LB strep) x 100. The efficiency of transformation was calculated as the percent of mutant transformed/average percent wild-type transformed.

### Statistical Analysis

GraphPad Prism software version 11 was used for all statistical analyses. Data was checked for normality with Shapiro-Wilk test. Parametric data were assessed with a one-sample Student’s t-test, Welch’s t-test, or Welch’s ANOVA post-hoc Dunnett’s test. Non-parametric data were analyzed with the Wilcoxon signed-rank test, the Kruskal-Wallis test with Dunn’s multiple comparisons, or the Mann-Whitney U test. Significance was set at p<0.05.

